# Evaluation of the Abbott ARCHITECT HIV Ag/Ab Combo Assay for Determining Recent HIV-1 Infection

**DOI:** 10.1101/2020.11.09.374017

**Authors:** Kelly A. Curtis, Donna L. Rudolph, Yi Pan, Kevin Delaney, Kathryn Anastos, Jack DeHovitz, Seble G. Kassaye, Carl V. Hanson, Audrey L. French, Elizabeth Golub, Adaora A. Adimora, Igho Ofotokun, Hector Bolivar, Mirjam-Colette Kempf, Philip J. Peters, William M. Switzer

**Affiliations:** Division of HIV/ AIDS Prevention, National Center for HIV/AIDS, Hepatitis, STD, and TB Prevention, Centers for Disease Control and Prevention, Atlanta, Georgia, United States of America; Women’s Interagency HIV Study (WIHS)

## Abstract

**Background:** Given the challenges and costs associated with implementing HIV-1 incidence assay testing, there is great interest in evaluating the use of commercial HIV diagnostic tests for determining recent HIV infection. A diagnostic test with the capability of providing reliable data for the determination of recent HIV infection without substantial modifications to the test protocol would have a significant impact on HIV surveillance. The Abbott ARCHITECT HIV Ag/Ab Combo Assay is an antigen/antibody immunoassay, which meets the criteria as the first screening test in the recommended HIV laboratory diagnostic algorithm for the United States.

**Methods:** In this study, we evaluated the performance characteristics of the ARCHITECT HIV Ag/Ab Combo signal-to-cutoff ratio (S/Co) for determining recent infection, including estimation of the mean duration of recent infection (MDRI) and false recent rate (FRR), and selection of recency cutoffs.

**Results:** The MDRI estimates for the S/Co recency cutoff of 400 is within the 4 to 12 months range recommended for HIV incidence assays, and the FRR rate for this cutoff was 1.5%. Additionally, ARCHITECT Combo S/Co values were compared relative to diagnostic test results from two prior prospective HIV-1 diagnostic studies in order to validate the use of the S/Co for both diagnostic and recency determination.

**Conclusion:** Dual-use of the ARCHITECT Combo assay data for diagnostic and incidence purposes would reduce the need for separate HIV incidence testing and allow for monitoring of recent infection for incidence estimation and other public health applications.

## Introduction

In 2014, Centers for Disease Control and Prevention (CDC) and the Association of Public Health Laboratories (APHL) issued updated HIV testing recommendations for laboratory diagnosis of HIV in the United States (US)[1]. The revised guidelines recommend use of Food and Drug Administration (FDA)-approved HIV tests with improved detection of acute HIV-1 infection, as well as HIV-2. The recommended diagnostic algorithm involves screening with a HIV-1/HIV-2 antigen (Ag)/antibody (Ab) combination immunoassay, followed by confirmation with an HIV-1/HIV-2 antibody differentiation immunoassay. Specimens that yield discordant or indeterminate immunoassay test results should be resolved with an HIV-1 nucleic acid test (NAT) to diagnose potential acute infection. Several studies have addressed the performance of the diagnostic algorithm with various combinations of FDA-approved diagnostic tests that meet the algorithm criteria [2–6]. The ARCHITECT HIV Ag/Ab Combo assay ([ARCHITECT] Abbott Laboratories, Chicago, IL) is one example of an HIV Ag/Ab combination immunoassay that is commercially available in the US and FDA-approved for HIV diagnosis.

The performance characteristics of the ARCHITECT suggest that the assay may also be useful for distinguishing recent from late or chronic HIV infection for the purposes of estimating HIV-1 incidence. The ARCHITECT is a chemiluminescent microparticle immunoassay (CMIA) that detects HIV-1/2 antibodies in serum or plasma, as well as HIV-1 p24 antigen. The chemiluminescent reaction resulting from the detection of HIV antibody and antigen is measured as relative light units (RLU) and a signal to cutoff ratio (S/Co) is calculated based on the reactivity of the specimen relative to an internal assay calibrator. Studies have demonstrated the ability of the ARCHITECT to detect acute infection, defined as HIV-1 NAT reactive and HIV-1 antibody negative [7–9], which is attributable to the sensitive detection of p24 antigen [10]. The ARCHITECT has a broad dynamic range for detection of the analytes and an association between S/Co and duration of infection has been demonstrated [11, 12].

HIV surveillance involves the collection of information related to new and existing cases of HIV infection and provides estimates of prevalence in a given population. The duration of HIV infection, however, cannot typically be inferred from routinely collected surveillance data and, therefore, estimation of incidence, the occurrence of new infections in a population, presents distinct challenges. Laboratory assays, developed or optimized for distinguishing recent from long-term HIV infection, allow for estimation of incidence based on a cross-sectional sampling of a population [13, 14]. Multiple HIV incidence assays have been evaluated for the measurement of a given biomarker, typically HIV antibody titers or avidity, including the commercially available HIV-1 LAg-Avidity EIA (Sedia Biosciences Corp., Portland, OR)[13, 15]. Testing for recent HIV-1 infection is typically independent of diagnostic testing, incurring additional costs, training needs, and dedicated equipment requirements. With the advent of new diagnostic technologies, there has been growing interest in evaluating select HIV diagnostic assays for determining recent infection.

In this study, we evaluated the feasibility of using the ARCHITECT S/Co to determine recent infection, using the standard assay protocol. The performance characteristics of the assay, such as the mean duration of recent infection (MDRI) and false recent rate (FRR), were estimated based on the S/Co from well-characterized subtype B HIV-1 seroconversion panels and optimal recency cutoffs were selected. The MDRI is defined as the average time that an HIV-infected person will spend in the “recent” state, as measured by a given incidence assay, while the FRR is a measure of the misclassification of long-term HIV infections as recent. HIV-1 recency status, based on the ARCHITECT S/Co values obtained through US surveillance systems, was compared to diagnostic test results from two prior prospective HIV-1 diagnostic studies.

## Materials and methods

### Seroconversion Panels and Long-term Specimens

For estimation of the ARCHITECT S/Co MDRI, 198 specimens from 26 antiretroviral therapy (ART)-naïve, subtype B HIV-1-infected subjects were evaluated (Table 1). Five HIV-1 seroconversion panels (n=42 specimens) were purchased from Zeptometrix Corp. (Buffalo, NY) and three panels (n=14) were obtained from SeraCare Life Sciences, Inc. (Milford, MA). A total of nine longitudinal seroconversion panels (n=82 specimens) were obtained through the Seroconversion Incidence Panel Project (SIPP) in collaboration with SeraCare Life Sciences, Inc., described in detail previously [16]. Lastly, nine recent seroconverters (n=60 specimens) were obtained from the Women’s Interagency HIV Study (WIHS). WIHS is an ongoing, prospective, multi-center cohort study aimed towards understanding disease progression in HIV-infected women (https://statepi.jhsph.edu/wihs/wordpress/)[17, 18]. WIHS was established in 1993 and recruited high-risk HIV-negative or HIV-infected women into ten clinical sites: Brooklyn, NY; Bronx, NY; Chicago, IL; Los Angeles, CA; San Francisco, CA; Washington, DC; Atlanta, GA; Birmingham, AL/ Jackson, MS; Chapel Hill, NC; Miami, FL. Each site obtained approval from the local Institutional Review Board (IRB) and participants provided written informed consent.

**Table 1.**
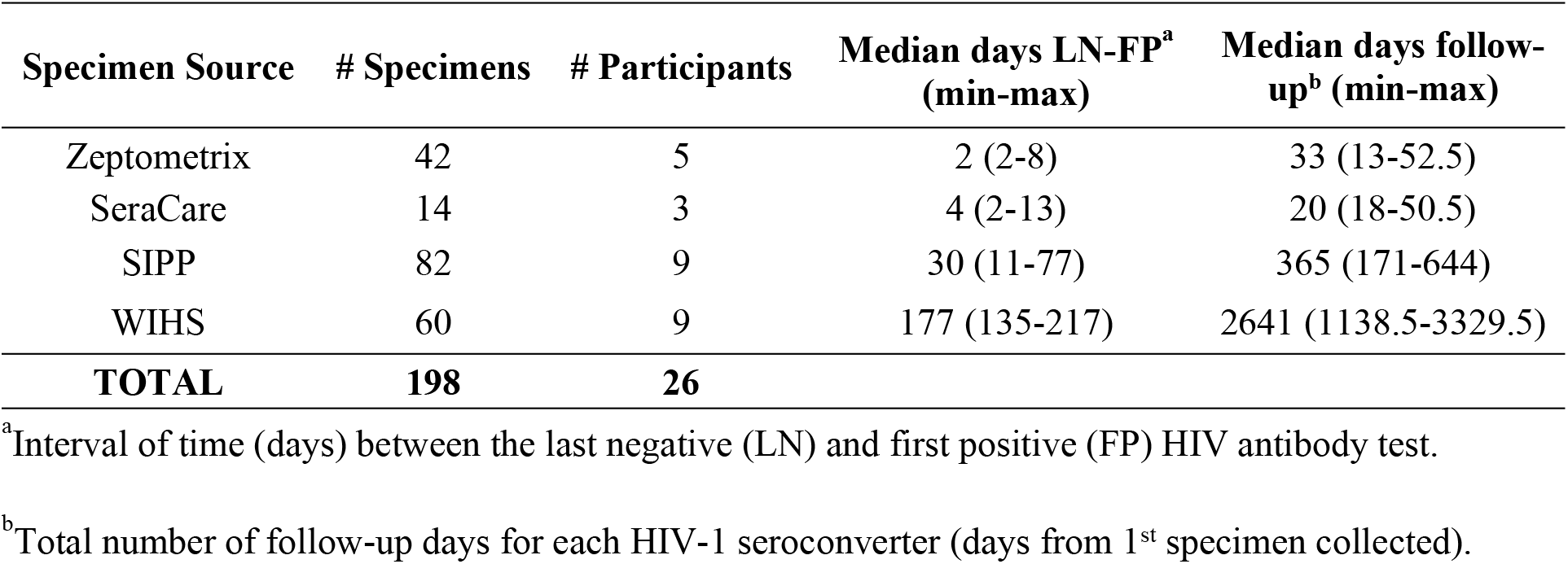
HIV-1 seroconversion panel characteristics.

| Specimen Source | # Specimens | # Participants | Median days LN-FP<sup>a</sup><br>(min-max) | Median days follow-up<sup>b</sup> (min-max) |
| --- | --- | --- | --- | --- |
| Zeptometrix | 42 | 5 | 2 (2-8) | 33 (13-52.5) |
| SeraCare | 14 | 3 | 4 (2-13) | 20 (18-50.5) |
| SIPP | 82 | 9 | 30 (11-77) | 365 (171-644) |
| WIHS | 60 | 9 | 177 (135-217) | 2641 (1138.5-3329.5) |
| **TOTAL** | **198** | **26** |  |  |

<sup>a</sup>Interval of time (days) between the last negative (LN) and first positive (FP) HIV antibody test.
<sup>b</sup>Total number of follow-up days for each HIV-1 seroconverter (days from 1<sup>st</sup> specimen collected).

For estimation of FRR, a total of 66 plasma specimens (single time points per study participant) were obtained from a 1982-1983 study in Atlanta, GA, involving subtype B HIV-1-infected men who have sex with men (MSM) diagnosed with lymphadenopathy [19–21]. The majority of the sample collection for the study occurred at a time prior to the advent of effective ART. The specimens included in this study were collected >2 years after the study entry date, since diagnostic test dates were not available for this cohort. All samples included in this study were unlinked from personal identifiers. CDC reviewed the protocol for the study and determined that CDC was not engaged in human subject research.

### ARCHITECT HIV Ag/Ab Combo Assay

The ARCHITECT HIV Ag/Ab Combo assay was performed according to the manufacturer’s instructions without modification. Plasma specimens were centrifuged for ten minutes at 8,000 rpm, transferred to polystyrene sample cups (Abbott Laboratories) in a total volume of 200μL, and then loaded onto the ARCHITECT i2000SR. The specimens were tested in singlicate due to volume limitations. The chemiluminescent signal, as measured by the instrument, is reported as a S/Co ratio of the relative light units (RLU) of the test sample to the RLU of the cutoff determined from an ARCHITECT calibrator. In addition to the kit controls, high and low external HIV-1 positive controls were included in each run. The controls were obtained from a proficiency testing panel derived from HIV-1-seropositive serum [22]. The high and low positive controls are consistent with long-term and recent infection, respectively, as measured by a previously characterized HIV incidence assay [23].

### Estimation of Mean Duration of Recent Infection (MDRI)

The MDRI was estimated with a linear binomial regression of a logit link with a cubic polynomial for the time since seroconversion for the probability of testing “recent” [24]. The 95% confidence interval was obtained through bootstrapping (by resampling subjects) with 10,000 bootstrap samples. Furthermore, for seroconversion panels from Zeptometrix Corp, SIPP and SeraCare Life Sciences, Inc., the subject’s testing history was used to obtain the estimated (earliest) date of detectable infection (EDDI). However, for the WIHS cohort, the seroconversion time was estimated as the middle point between the last negative and first positive test date due to lack of testing history data. Data points longer than 800 days from the last negative and the first positive were excluded from the analysis. The R package “inctool” was used for MDRI estimation (ref: https://cran.r-project.org/web/packages/inctools/index.html). The MDRI was estimated at ARCHITECT S/Co values of 300, 350, 400, 450, 500, 550, and 600.

### Estimation of False-recent Rate (FRR)

The FRR is the probability that a subject, who is infected longer than T (defined time after infection), will produce a “recent” infection result [15, 24]. In our study, samples used to calculate the false-recent rate were from people with HIV infection for greater than T, defined as 730.5 days or two years. The recency cutoffs of 300, 350, 400, 450, 500, 550 and 600 were used for estimating FRR. The FRR was estimated with a random sampling of the MSM cohort (N=66) based on availability of sufficient sample volume (Table 2).

**Table 2.** ARCHITECT HIV Ag/Ab Combo S/Co MDRI and FRR estimates.

| **Cutoff** | **MDRI<sup>a</sup> (95% CI)** | **N recent** | **FRR MSM<sup>b</sup> (95% CI)** |
| --- | --- | --- | --- |
| 300 | 103 (86, 130) | 74 | 0.0 (0.0, 0.05) |
| 350 | 145 (101, 224) | 80 | 0.0 (0.0, 0.05) |
| 400 | 190 (120, 281) | 88 | 1.5 (0.0004, 0.08) |
| 450 | 247 (145, 356) | 94 | 4.6 (0.01, 12.7) |
| 500 | 291 (179, 406) | 102 | 7.6 (2.5, 16.8) |
| 550 | 366 (222, 483) | 114 | 7.6 (2.5, 16.8) |
| 600 | 445 (283, 564) | 125 | 9.1 (3.4, 18.7) |

<sup>a</sup>N= 198 specimens (26 Subjects)
<sup>b</sup>N = 66 specimens (66 subjects)
MDRI=mean duration of recent infection (days); FRR=false recent rate (%); CI=confidence interval

### Comparison of Recency and Diagnostic Test Results

In this study, 681 specimens from the Screening Targeted Populations to Interrupt On-going Chains of HIV Transmission with Enhanced Partner Notification (STOP, n = 395)[25] and Los Angeles Rapid Test (LA Study, n = 286)[9, 26, 27] studies with available ARCHITECT, Multispot or Geenius, and Aptima Qualitative assay or HIV-1 viral load (VL) results were evaluated to assess the use of ARCHITECT for determining recent infection at each selected S/Co recency cutoff. Additionally, 661 specimens (n =98, STOP; n =563, LA Study) from persons with established infection (both ART naïve and receiving ART) with HIV-1 VL data (COBAS AmpliPrep/COBAS TaqMan® HIV-1 Test, v2.0, Roche Molecular Diagnostics) were included to evaluate the impact of VL on the performance of the ARCHITECT.STOP was a prospective study evaluating the recommended diagnostic algorithm for detection of acute HIV infection and linkage to enhanced partner services in New York City, NY; San Francisco, CA; and Raleigh, Durham, and Winston-Salem, NC. The LA Study evaluated the performance characteristics of six rapid HIV tests and was conducted at two clinical sites: the Los Angeles Gay and Lesbian Center and the Altamed Clinic, which primarily serve persons at high risk for HIV infection [26].

### Data Analysis

Differences between various groups of ARCHITECT S/Co values was determined using the Mann-Whitney U test. For the purposes of this study, two acute groups were defined based on the sequence of HIV test results. Acute group 1 was defined as ARCHITECT negative, Multispot/Geenius negative or indeterminate, and Aptima/VL reactive. Acute group 2 was defined as ARCHITECT reactive, Multispot/Geenius negative or indeterminate, and Aptima/VL reactive. Established infections were defined as having a reactive test result for all three tests.

The number of recent infections for the STOP and LA studies was determined based on the number of persons with ARCHITECT S/Co values below each of the evaluated recency cutoffs, with the exception of 300 and 350. These two recency cutoffs were not evaluated due to the extremely short MDRI estimates.

## Results

### Mean Duration of Recent Infection and False Recent Rate

The ARCHITECT S/Co for 198 specimens from 26 recent HIV-1 seroconverters demonstrated a rapid increase in assay values within the first year post-seroconversion, allowing for a clear distinction to be made between recent (<6 months) and long-term (> 2 years) infection (Fig 1A). The separation in S/Co values was further illustrated through the direct comparison of S/Co values at <6 months to >6 months post-seroconversion (Fig 1B). No overlap was observed in the 25^th^ to 75^th^ percentile of reactivity and higher mean values were noted at >6 months as compared to <6 months (P<0.0001). Furthermore, a difference in S/Co values (P<0.0001) was also observed between the recent group (<6 months post-seroconversion) compared to values for the 248 persons with long-term infection (>2 years post-seroconversion). The mean S/Co was 228.3, 726.9, and 731.0 for the <6 months, > 6 months, and >2 year post-seroconversion groups, respectively.

**Fig 1.**
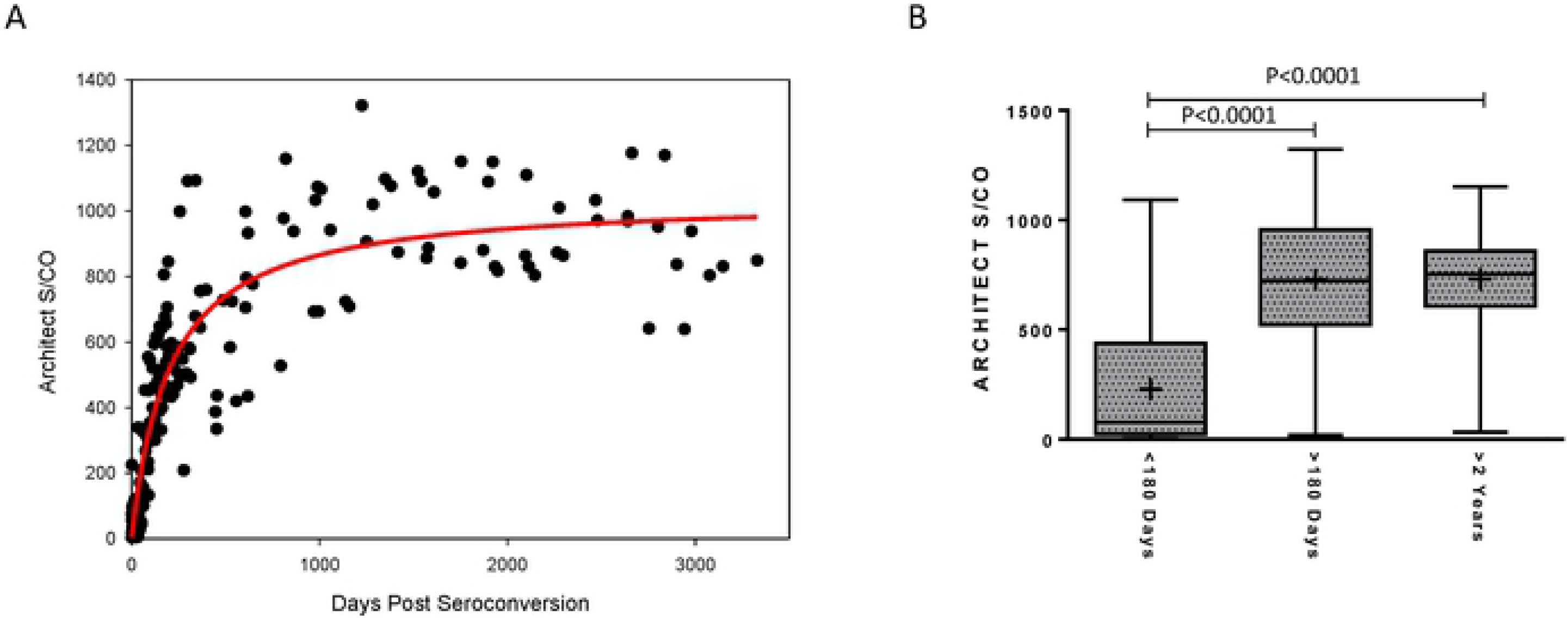
Longitudinal ARCHITECT HIV Ag/Ab Combo signal-to-cutoff (S/Co) values from recent HIV-1 seroconverters. **(A)** The S/Co ratios for longitudinally collected specimens from 26 antiretroviral therapy-naive recent seroconverters (N=198) were plotted over days since estimated seroconversion. The solid red line represents the logarithmic curve fit to the data using non-linear regression. **(B)** Box plots show the 25th to 75th percentile of the S/Co ratios at indicated days post-seroconversion, while the middle lines represent the median values and whiskers represent the minimum and maximum values. The mean is indicated by “+”.

The MDRI estimated at the ARCHITECT S/Co cutoffs of 300, 350, 400, 450, 500, 550, and 600 are summarized in Table 2. The MDRI ranged from a minimum of 103 to a maximum of 445 days, increasing with higher cutoff values. The MDRI at cutoffs 400, 450, 500, and 550 were within six months to one year, at 190, 247, 291, and 366 days, respectively. The FRR for the MSM cohort, ranged from 0.0 to 9.1% at the different S/Co cutoffs evaluated.

### Comparison of HIV-1 Recency and Diagnostic Test Results

The number of HIV diagnoses from both the STOP and LA studies is summarized by category (acute group 1, acute group 2, and established) in Table 3. For both studies, the majority of diagnoses (77%) fell within the established category (64% STOP, 94% LA study). For both acute groups, the number of recent infections was highly consistent for all cutoffs evaluated. All samples from persons in acute group 1 were ARCHITECT negative, as samples from these individuals were HIV-1 NAT reactive only and determined to be from persons with acute infection based on the CDC/APHL HIV laboratory testing algorithm. A total of 132 out of 133 samples (99.2%) in acute group 2 were determined to be from persons with recent infection (131 out of 133 for the lowest cutoff of 400). The number of recent infections identified in the established group was variable, ranging from 54% to 80% of the total established infections, increasing with higher recency cutoff value.

**Table 3.**
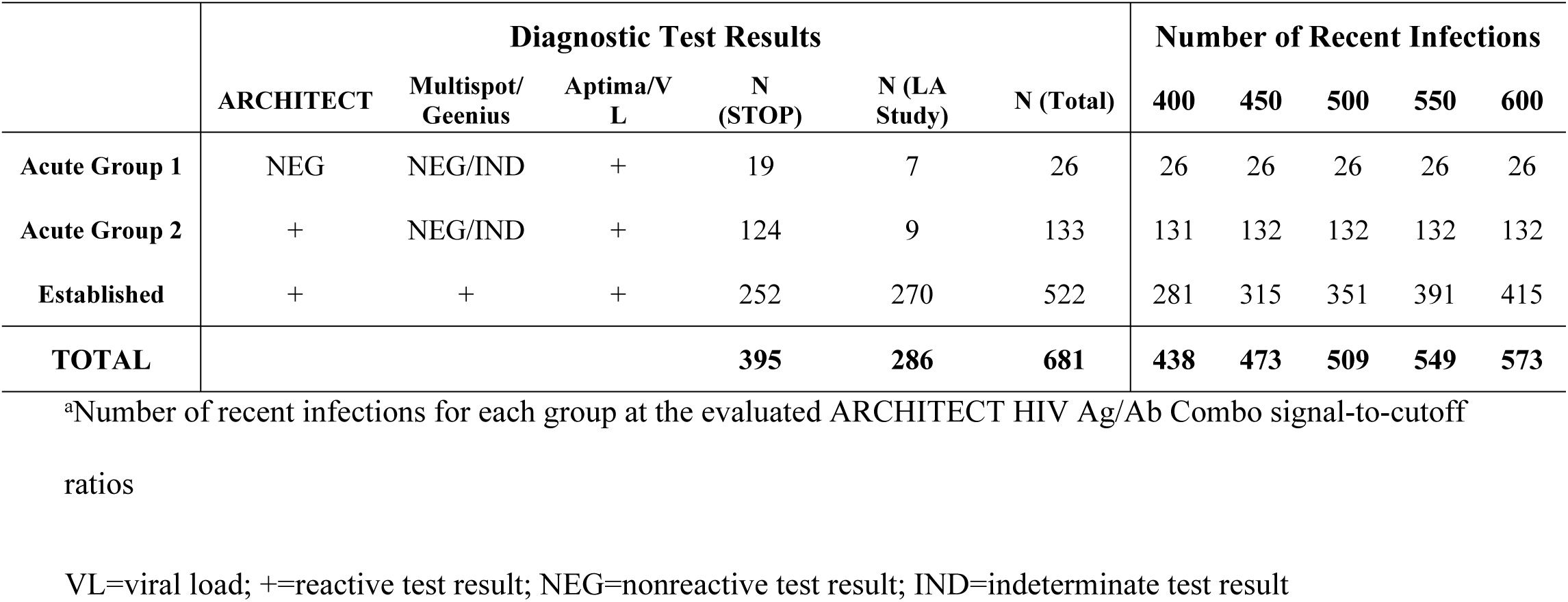
HIV diagnostic test results summary for STOP and LA study cohorts.

The distribution of ARCHITECT S/Co for the acute and established groups is shown in Fig 2A. The S/Co for all specimens within acute group 1 is below the limit of detection for the ARCHITECT. The mean S/Co was 60.4 and 407.8 for specimens from acute group 2 and the established group, respectively. A difference (P<0.0001) was noted between the S/Co values for acute group 2 and the established group. ARCHITECT S/Co values for the established groups from the STOP and LA studies were further evaluated based on VL and reported ART use (Fig 2B). Higher S/Co values (P=0.0472) were observed in the established group with a VL > 1,000 RNA copies/mL compared to a VL < 1,000 RNA copies/mL (median S/Co of 422.7 versus 394.9, respectively). Interestingly, a higher median S/Co ratio (P=0.0002) was observed in the ART-use group compared to the ART-naive group (median S/Co of 419.7 versus 373.8, respectively).

**Figure 2.**
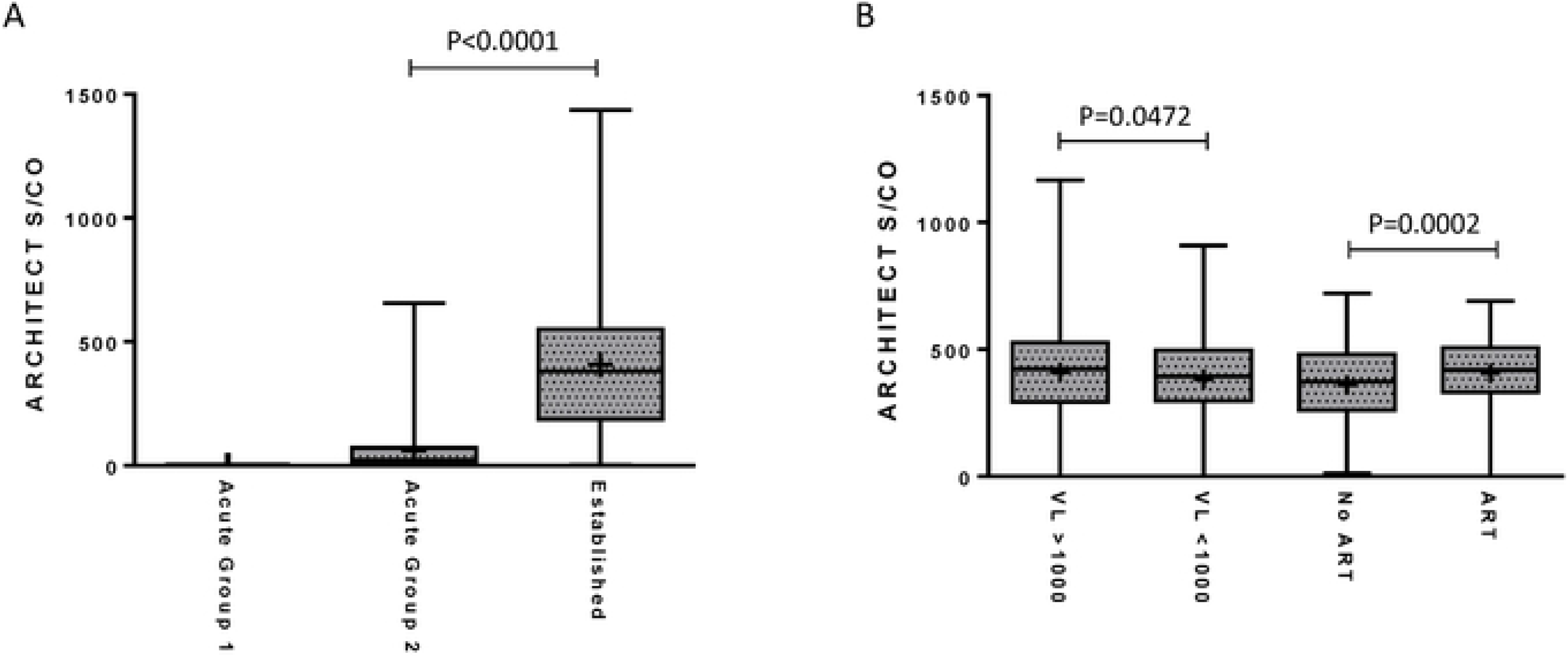
Performance of the ARCHITECT HIV Combo Assay signal-to-cutoff (S/Co) in Determining Recent Infection. Box plots comparing the ARCHITECT HIV Ag/Ab Combo S/Co ratios in the acute and established groups **(A)**, the low (<1,000 RNA copies/mL; N=298) versus high (>1,000 RNA copies/mL; N=363) plasma viral loads (VL) samples, and no antiretroviral therapy (ART; N=370) versus ART-use groups (N=276) **(B)** in the STOP and LA study cohorts. The boxes represent the 25th to 75th percentile of the S/Co ratios, while the middle lines represent the median values and whiskers represent the minimum and maximum values. The mean is indicated by “+”.

## Discussion

Given that HIV incidence is a valuable measure for monitoring HIV trends over time in a population, there is a continued goal to develop and evaluate methods that would reduce error associated with these estimates and to expand HIV incidence testing in settings where additional laboratory testing is not feasible. In this study, we demonstrated the performance of the ARCHITECT, an HIV-1/HIV-2 antigen/antibody combination immunoassay, for determining recent infection. The data presented in this study suggest that S/Co values obtained with the ARCHITECT can serve a dual purpose, providing both HIV diagnostic test results as well as recency data that can be incorporated into population-level incidence estimates or used for other research purposes, requiring no additional testing or modifications to the assay protocol. A S/Co recency cutoff of 400 meets the recommended acceptance criteria for HIV incidence assays for evaluation of subtype B HIV-1 incidence. Since ARCHITECT S/Co values are available through diagnostic screening, the costs associated with estimating HIV incidence would be significantly reduced, potentially expanding the capability of some laboratories to provide recency data for population-based incidence estimation.

Recently, Suligoi *et al*. performed an avidity modification with the ARCHITECT and demonstrated that an avidity index, which is a measure of the binding strength between antibody and antigen, is an accurate marker for discriminating recent from long-term infection [28]. In this study, we showed that the ARCHITECT S/Co values, alone, from longitudinal seroconversion panels exhibited a biomarker maturation curve (Fig 2A) similar to other well-performing or promising incidence assays [23, 29], which is suggestive of an assay’s ability to effectively distinguish recent from long-term infection. Similarly, Grebe *et al.* demonstrated comparable performance between the ARCHITECT S/Co using an unmodified protocol and HIV-1 LAg-Avidity EIA [12]. However, performance characteristics of the ARCHITECT for determining recency cannot be compared between the two studies given the notable differences in subtype diversity and inclusion criteria for the specimens used to estimate MDRI. The performance of the Geenius HIV-1/2 Supplemental Assay, a single-use immunochromatographic assay that detects and differentiates antibodies to HIV-1 and HIV-2, was also evaluated for determining recent infection and preliminary findings indicate that the assay may be useful for both HIV diagnosis and recency determination [24]. One caveat to this approach is that recency is determined for the Geenius assay based on band intensity; however, the raw band intensity values are not readily accessible through the Geenius reader for the standard user limiting its utility. Given that the ARCHITECT and Geenius are frequently used concurrently for diagnosis of HIV in the US, an algorithm composed of both test results may be considered for determining recent HIV infection.

ARCHITECT S/Co values are routinely collected in some US laboratories through the implementation of the HIV diagnostic testing algorithm; therefore, we were able to demonstrate the relationship between the HIV diagnostics test results and recency determination for two prospective diagnostic studies. Here, we determined recency at the individual level, though HIV recency data are predominantly used for estimation of population-level incidence. The benefits of recency determination at the individual level are yet to be fully explored; however, there may be utility in capturing the number of recent infections in a cohort for research purposes. As demonstrated with the STOP and LA cohorts, new diagnoses determined to be established infections by the diagnostic testing algorithm include both long-term infections, as well as persons within the MDRI of the recency assay, as HIV antibody has exceeded the limit of detection for diagnostic tests but antibody titers are continuing to increase or evolve. Overall, these data suggest that the ARCHITECT S/Co ratio is predictive of time since seroconversion and has the capability to reliably detect changes in biomarker levels over time.

Although the performance characteristics of the ARCHITECT presented here are promising, additional data are needed to fine-tune the MDRI and FRR estimates. A limitation of this study is the relatively small number and limited diversity of specimens available for the MDRI and FRR analyses. Certain population characteristics, such as ART use, may greatly impact FRR estimates. The MSM cohort evaluated in this study is unique in that the original study enrolled participants at a time prior to the availability of effective ART, so ART likely had little to no impact on the FRR estimates. Furthermore, Abbott has recently introduced the Alinity family of diagnostic testing platforms. Further investigation is needed to determine whether the ARCHITECT and Alinity platforms can be used interchangeably for determining recent HIV infection, though an initial evaluation has indicated comparable performance for detection of HIV [30]. ART-use has proven to be problematic for most HIV incidence assays and a slight, yet significant, reduction in the median S/Co in our study was associated with VLs less than 1,000 HIV RNA copies/mL for the ARCHITECT. However, sufficient data were not available to determine whether ART-induced virus suppression leads to increased FRRs. Prior studies have recommended that HIV incidence assays be used in conjunction with VL to eliminate potential false-recent results due to suppressed VLs in persons receiving ART and elite controllers [13, 31].

In summary, the data presented here suggest that the performance characteristics of the ARCHITECT HIV Ag/Ab Combo test meets the acceptance criteria for a HIV incidence assay and, for laboratories that employ the platform for HIV diagnostic testing, the utility of the assay may be expanded for additional surveillance applications.

## Acknowledgements

The authors would like to thank Alexandra Tejada at the CDC for providing training and assistance with the ARCHITECT platform. The authors would also like to thank Eduard Grebe for valuable discussions on the MDRI estimation. Use of trade names is for identification only and does not imply endorsement by the U.S. Department of Health and Human Services, the Public Health Service, the National Institutes of Health (NIH), or the CDC. The findings and conclusions in this report are those of the authors and do not necessarily represent the views of the NIH or CDC.

Data in this manuscript were collected by the Women’s Interagency HIV Study (WIHS). WIHS (Principal Investigators): UAB-MS WIHS (Mirjam-Colette Kempf and Deborah Konkle-Parker), U01-AI-103401; Atlanta WIHS (Ighovwerha Ofotokun and Gina Wingood), U01-AI-103408; Bronx WIHS (Kathryn Anastos and Anjali Sharma), U01-AI-035004; Brooklyn WIHS (Howard Minkoff and Deborah Gustafson), U01-AI-031834; Chicago WIHS (Mardge Cohen and Audrey French), U01-AI-034993; Metropolitan Washington WIHS (Seble Kassaye), U01-AI-034994; Miami WIHS (Margaret Fischl and Lisa Metsch), U01-AI-103397; UNC WIHS (Adaora Adimora), U01-AI-103390; Connie Wofsy Women’s HIV Study, Northern California (Ruth Greenblatt, Bradley Aouizerat, and Phyllis Tien), U01-AI-034989; WIHS Data Management and Analysis Center (Stephen Gange and Elizabeth Golub), U01-AI-042590; Southern California WIHS (Joel Milam), U01-HD-032632 (WIHS I – WIHS IV). The WIHS is funded primarily by the National Institute of Allergy and Infectious Diseases (NIAID), with additional co-funding from the Eunice Kennedy Shriver National Institute of Child Health and Human Development (NICHD), the National Cancer Institute (NCI), the National Institute on Drug Abuse (NIDA), and the National Institute on Mental Health (NIMH). Targeted supplemental funding for specific projects is also provided by the National Institute of Dental and Craniofacial Research (NIDCR), the National Institute on Alcohol Abuse and Alcoholism (NIAAA), the National Institute on Deafness and other Communication Disorders (NIDCD), and the NIH Office of Research on Women’s Health. WIHS data collection is also supported by UL1-TR000004 (UCSF CTSA), UL1-TR000454 (Atlanta CTSA), and P30-AI-050410 (UNC CFAR).

The STOP study was supported by a cooperative agreement between the Centers for Disease Control and Prevention (CDC) and the San Francisco Department of Public Health (5U01PS001564), New York City Department of Health and Mental Hygiene (5U01PS001561), and the University of North Carolina at Chapel Hill (5U01PS001559).

The LA Rapid testing study was supported through a cooperative agreement with the Health Research Association of Los Angeles County (U64 CCU919524). The authors would like to thank the study participants and staff, especially Apurva Uniyal, Peter Kerndt, Lashawnda Royal, and Staeci Morita.

## Notes

**Conflicts of Interest and Sources of Funding:** No conflicts of interest were declared. This work was supported through intramural CDC funding.

